# Optimizing single-cell RNA sequencing methods for human colon biopsies: droplet-based vs. picowell-based platforms

**DOI:** 10.1101/2024.06.24.600526

**Authors:** Jonathan M. Downie, Ryan J. Musich, Connor M. Geraghty, Alexander Caraballo, Shijie He, Saleh Khawaled, Kylor Lachut, Timothy Long, Julie Y. Zhou, Omer H. Yilmaz, Thaddeus Stappenbeck, Andrew T. Chan, David A. Drew

**Affiliations:** Clinical and Translational Epidemiology Unit, Harvard Medical School/Massachusetts General Hospital, Boston, MA; Division of Gastroenterology, Massachusetts General Hospital, Boston, MA; Inflammation and Immunity, Cleveland Clinic, Cleveland, OH; Koch Institute for Integrative Cancer Research, Massachusetts Institute of Technology, Cambridge, MA

**Keywords:** Single-cell transcriptomics, human colorectal biopsies, mucosal biology, epithelial cells

## Abstract

**Background & Aims:** Single-cell RNA sequencing (scRNA) has empowered many insights into gastrointestinal microenvironments. However, profiling human biopsies using droplet-based scRNA (D-scRNA) is challenging since it requires immediate processing to minimize epithelial cell damage. In contrast, picowell-based (P-scRNA) platforms permit short-term frozen storage before sequencing. We compared P- and D-scRNA platforms on cells derived from human colon biopsies.

**Methods:** Endoscopic rectosigmoid mucosal biopsies were obtained from two adults and conducted D-scRNA (10X Chromium) and P-scRNA (Honeycomb HIVE) in parallel using an individual’s pool of single cells (> 10,000 cells/participant). Three experiments were performed to evaluate 1) P-scRNA with cells under specific storage conditions (immediately processed [fresh], vs. frozen at -20C vs. -80C [2 weeks]); 2) fresh P-scRNA versus fresh D-scRNA; and 3) P-scRNA stored at -80C with fresh D-scRNA.

**Results:** Significant recovery of loaded cells was achieved for fresh (80.9%) and -80C (48.5%) P-scRNA and D-scRNA (76.6%), but not -20C P-scRNA (3.7%). However, D-scRNA captures more typeable cells among recovered cells (71.5% vs. 15.8% Fresh and 18.4% -80C P-scRNA), and these cells exhibit higher gene coverage at the expense of higher mitochondrial read fractions across most cell types. Cells profiled using D-scRNA demonstrated more consistent gene expression profiles among the same cell type than those profiled using P-scRNA. Significant intra-cell-type differences were observed in profiled gene classes across platforms.

**Conclusions:** Our results highlight non-overlapping advantages of P-scRNA and D-scRNA and underscore the need for innovation to enable high-fidelity capture of colonic epithelial cells. The platform-specific variation highlights the challenges of maintaining rigor and reproducibility across studies that use different platforms.

## Background

Single-cell RNA sequencing (scRNA) has significantly expanded our understanding of multiple cellular microenvironments, including the gastrointestinal (GI) mucosal microenvironment that broadly consists of immune, stromal, and epithelial cell subtypes. However, high-fidelity capture of epithelial cells from human biopsy samples for transcriptional profiling remains challenging [1]. Prior scRNA studies of patient-derived colon biopsies have primarily utilized droplet-based scRNA (D-scRNA) platforms, but have well-described high rates of mitochondrial contaminant reads per cell [2, 3], which is a signature of dying or stressed cells that can result in potentially aberrant transcriptional profiles [4]. This impacts epithelial cells in particular since they undergo programmed cell death upon detachment from the basement membrane [5], which occurs during tissue disassociation necessary for scRNA sequencing. As several GI conditions, particularly the formation of neoplasia, originate from the intestinal epithelium [6], data loss within these cell types can significantly impact experiments focusing on mucosal biology and disease processes. The observed loss of cellular integrity is likely partly due to the characteristic fragility of these cell types once dissociated into single-cells *ex vivo* and exacerbated by the limitations associated with requiring fresh tissue. The time from fresh biopsy to single-cell library preparation can be variably impacted by the proximity of clinical areas to processing laboratories, procedural delays, and the availability of resources for immediate library preparation, leading to significant noise in D-scRNA datasets, even before considering heterogeneity introduced via biological or individual variability.

Some of these limitations may be overcome with the optimization of fresh tissue procurement and processing or with combinatorial approaches using fixed cell approaches (i.e., ATAC-seq or combinatorial barcoding) for epithelial cell profiling and droplet-based approaches for the remaining compartments; however, this is less efficient, cost-effective, and integrated than using a single approach. In contrast to D-scRNA, picowell scRNA (P-scRNA) systems partition single cells within microwells containing library preparation agents. Some P-scRNA systems are designed to better capture fragile cell types related to abbreviated processing approaches [7]. They also provide an opportunity for short-term cryopreservation to batch library preparation/sequencing to reduce overall costs and improve access/portability. However, P-scRNA systems may capture fewer cells per experimental setup compared to D-scRNA, thereby potentially limiting the profiling of rarer cell types. To more fully assess the relative advantages and disadvantages of each platform, we compared data from cells derived from the same human colon biopsy specimens that were profiled through D-scRNA and P-scRNA, including P-scRNA under different storage conditions.

## Results

Rectosigmoid colon mucosal biopsies were obtained during flexible sigmoidoscopy of two adults (participants #1 and #2; 6 biopsies each). The biopsies were immediately dissociated into participant-specific pools of single-cells, (**Fig. 1A, Methods**) flow-sorted, and split for D-scRNA (10X Chromium) and P-scRNA (Honeycomb HIVE) library preparation in parallel, according to the manufacturer’s standard protocols. We performed three primary experiments using these data to compare the effectiveness of different storage conditions for P-scRNA and then performed D-scRNA and P-scRNA head-to-head comparisons (**Fig. 1A**). To perform these comparisons and reduce the potential for platform-related batch effects, we developed a pipeline using open-source tools to generate per cell gene counts the same manner for D-scRNA and P-scRNA libraries (**Methods)**. Among P-scRNA clusters, no clear transcriptional or batch effects between fresh samples and samples stored in frozen conditions were evident (**Fig. 1B, C**). However, stark platform-specific differences were evident between P-scRNA and D-scRNA libraries when visualizing data using unsupervised UMAP (**Fig. 1C**). After performing sample integration (**Methods**) to account for platform and storage conditions related to batch effects, UMAP projections demonstrate overlap in the single-cell populations captured by P-scRNA and D-scRNA (**Fig. 1D**).

**Figure 1.**
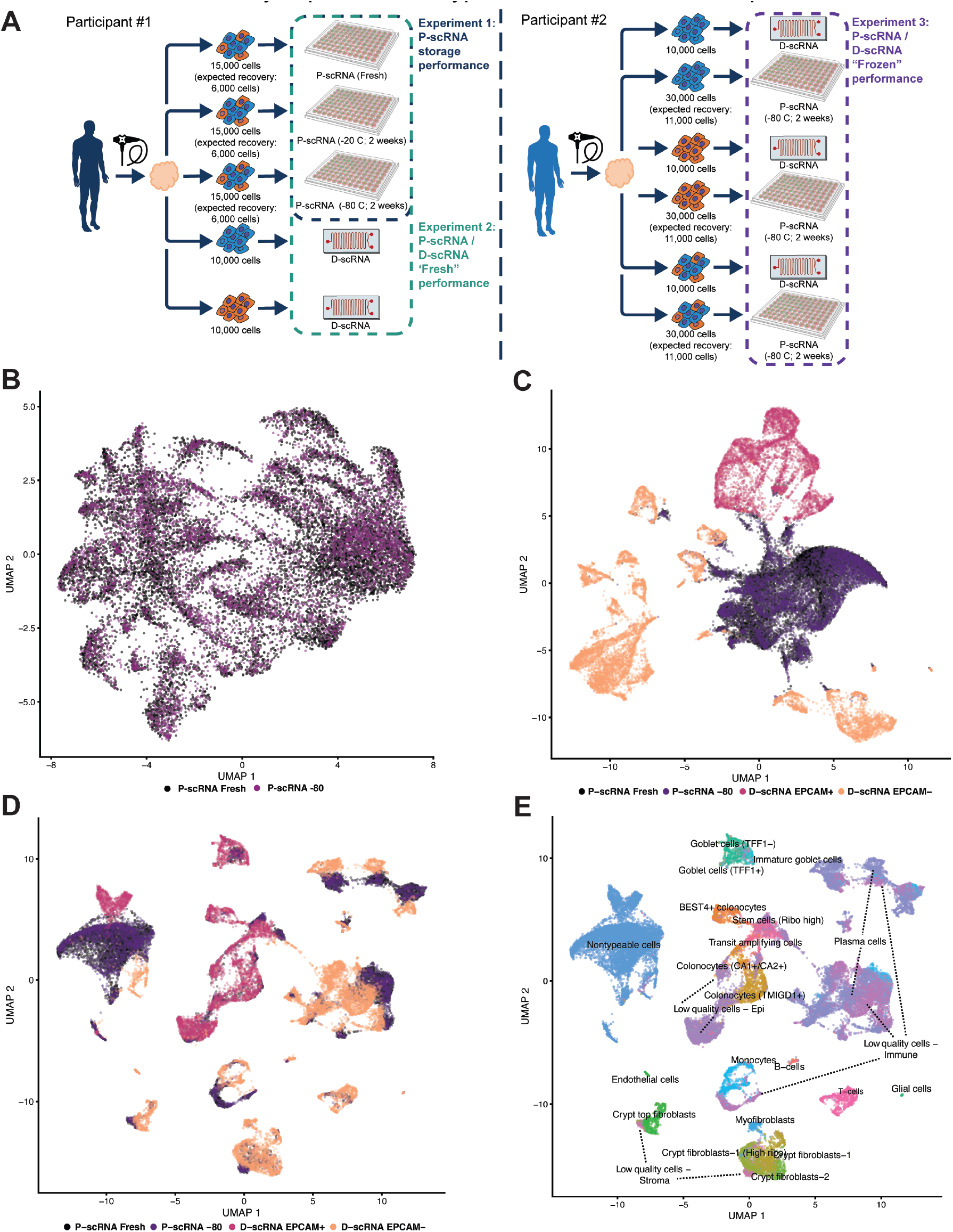
Overview of P-scRNA versus D-scRNA head-to-head differences. **A**) A schematic of the study design. Single-cell pools from two individual participants informed three experiments (Experiments 1-3) to compare i) P-scRNA fresh preparations vs. frozen storage conditions at -20C or -80C within Participant 1; ii) an initial comparison of P-scRNA vs D-scRNA results within Participant 1; and iii) a follow-up comparison of P-scRNA vs. D-scRNA results within Participant 2 that more stringently controlled for the amount of cells loaded for capture. **B-D**) UMAP projections of P-scRNA fresh vs. -80 frozen storage libraries from Participant 1 show similar transcriptional profiles (B), but P-scRNA vs. D-scRNA comparisons highlight significant global platform-specific effects **C)**. After adjusting for platform and storage conditions (**D and E**), projections demonstrate that both methods reasonably capture all cell types, with few notable exceptions.

### Experiment 1

Because the option to freeze cells prior to library preparation is a key feature of P-scRNA, we first assessed the impact of short-term (2 weeks) cryopreservation at -20C or -80C on P-scRNA compared to P-scRNA library preparation on fresh samples as required for the D-scRNA platform. Using the single-cell pool from *Participant 1*, we prepared three P-scRNA chambers each with live single cells combined at 1:1 with viable EPCAM+/CD45-“epithelial” cells and EPCAM- “non-epithelial” flow-sorted cells (n=7,500 of each cell type) to ensure balanced representation of cell compartments. According to the manufacturer, the loading of this total number of cells in each chamber (N=15,000) was expected to recover approximately 6,000 cells {https://honeycomb.bio/faqs/}. The first P-scRNA chamber was used to prepare P-scRNA libraries immediately (“Fresh”) and then sequenced, undergoing initial QC to remove doublets and cells with low gene counts, recovering data for 12,141 cells (80.9% of loaded cells). The second chamber was frozen for two weeks at -20C (manufacturer-recommended storage condition) and then underwent library preparation, sequencing, and identical QC procedures, recovering data from only 562 cells (3.7% recovered). The third chamber was frozen for two weeks at -80C and then underwent library preparation, sequencing, and QC, recovering data for 7,279 cells (48.5% recovered).

Due to the poor performance of the -20C storage condition, we focused all subsequent analyses on comparing cell clustering and typing on sequencing data derived from the chambers with cells prepared and sequenced immediately (fresh) with data derived from the chamber frozen for two weeks at -80C prior to preparation and sequencing. A large proportion of cells under both conditions formed a cluster (Fresh n=4,783; -80C n=2,531 cells) that expressed non-specific, multi-compartment markers (**Supplementary Fig. 1)**, suggesting the presence of multiplets not otherwise removed by our analysis pipeline which were removed manually *post hoc*. A large number of additional cells (Fresh n=5,438; -80C=3,409 cells) could be typed to a mucosal compartment – epithelial, immune, or stromal – but not with greater granularity due to low quality (**Supplementary Fig. 1)**, leaving 1,920 Fresh and 1,339 -80C cells, or 15.8% and 18.4% of recovered cells, in which a specific cell type could be inferred (**Fig. 2A**). This demonstrates comparable performance between the two conditions, but with significantly lower-than-expected yield under both conditions. Notably, however, most epithelial cell lineages, except for rare cell types such as enteroendocrine and tuft cells, were represented in both the Fresh and -80C libraries.

**Figure 2.**
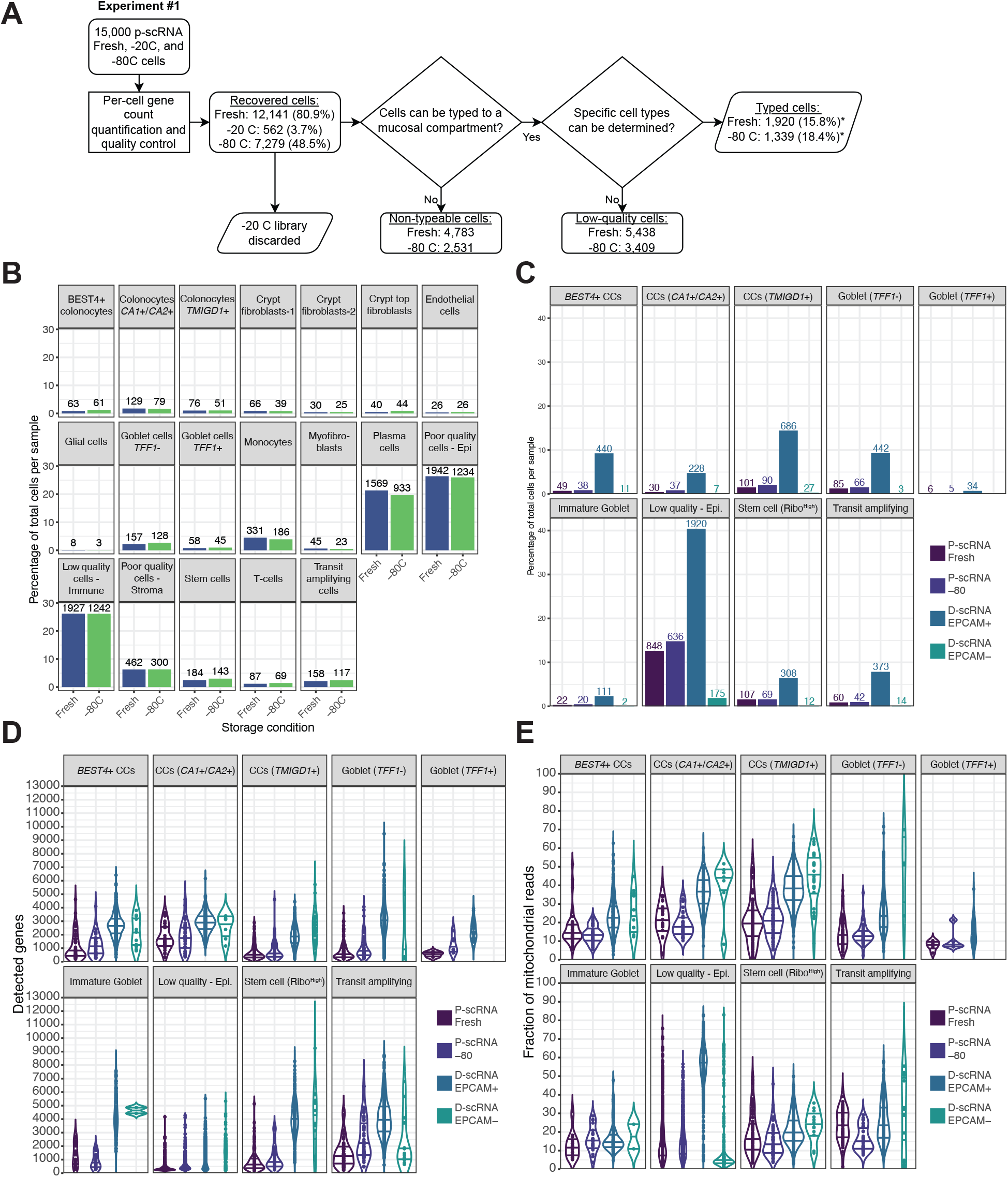
Comparisons of P-scRNA storage conditions versus D-scRNA (Experiments 1 & 2). **A**) Percentage of typable cell counts per cell type for the Fresh vs -80C P-scRNA comparison with labeled cell count numbers. **B**) Percentage of typable cell counts per epithelial cell type for the P-scRNA Fresh/-80 vs. D-scRNA comparison with labeled cell count numbers. **C**) Number of detected genes and **D**) mitochondrial read fraction per epithelial cell type for the P-scRNA Fresh/-80 vs. D-scRNA comparison with labeled cell count numbers. Horizontal lines within violin plots represent the 25^th^, 50^th^, and 75^th^ percentile levels. CC = colonocytes

### Experiment 2

To compare platform-specific differences, we prepared a D-scRNA library from an estimated 10,000 EPCAM+/CD45-“epithelial” sorted cells and another D-scRNA library from 10,000 EPCAM-“non-epithelial” sorted cells derived from the same single-cell pool (Participant 1) used in Experiment 1. As is required by D-scRNA libraries were immediately prepared from fresh samples. Sequenced libraries were jointly analyzed with P-scRNA sequenced libraries from Participant 1. Within the D-scRNA data, of the combined 20,000 cells loaded, 15,318 cells were recovered (76.6%), yielding comparable cell recovery to the freshly processed P-scRNA samples (81%). Notably, because the epithelial compartment (EPCAM+/CD45-) was prepared in parallel with the non-epithelial compartment (EPCAM-) for D-scRNA, stark differences in recovery were observed between compartments. Only 57.5% of epithelial cells (5,750 cells) were recovered compared to 95.7% of non-epithelial cells (9,568 cells), highlighting the challenge in recovering epithelial cells using D-scRNA. Among D-scRNA recovered cells, after the removal of non-typeable cells, 4,750 epithelial (EPCAM+/CD45-) and 9,255 non-epithelial (EPCAM-) remained. Of these, 41.4% and 30.0% were able to be typed to a mucosal compartment, but not with cell type specificity (i.e. low-quality cells) (**Figure 2B and Supplementary Fig. 2A**), resulting in 3,782 epithelial and 7,163 non-epithelial cells, or 65.8% and 74.9% of recovered cells, that could be assigned a specific cell-type. In addition, as a result of merging D-scRNA and P-scRNA sequenced libraries, some low-quality cells captured and previously mapped to a mucosal compartment were no longer able to be assigned to a mucosal compartment (e.g. 643 cells in the Fresh P-scRNA library), likely owing to the higher resolution data present in D-scRNA cell libraries driving tighter cluster assignments. Moreover, both D-scRNA libraries had higher unique molecular identifiers (UMIs) and detected gene counts across almost all cell types/compartments compared to Fresh P-scRNA (**Figure 2B-C, Supplementary Figs. 2B-C**), though higher mitochondrial DNA (mtDNA) read counts were observed among most D-scRNA cell types (**Figure 2D, Supplementary Fig. 2D)**. The trend of lower gene counts, UMI counts, and mtDNA read fractions present in the fresh P-scRNA sample was also present in the -80C P-scRNA library (**Supplementary Fig. 2**), suggesting this is a finding attributed to the platform, not a specific storage condition.

### Experiment 3

Based on these results establishing that P-scRNA libraries derived from fresh samples were similar to those cryopreserved at -80C for two weeks, we next sought to more closely compare data generated by D-scRNA and P-scRNA approaches as they would likely be employed (Fresh D-scRNA vs. -80C stored P-scRNA) while more tightly controlling for input cellular yields across platforms. From a second independent participant (Participant 2), we dissociated biopsies into a single cell pool and sorted them into three populations: 1) live “mixed” cells (sorted only for viability but not partitioned according to EPCAM/CD45 marker expression; 2) live “epithelial” cells (EPCAM+/CD45-), and 3) live “non-epithelial” cells (EPCAM-). From each of these populations, we split cells to generate three D-scRNA and three P-scRNA libraries. D-scRNA libraries were prepared immediately for each population (“Mixed”, “Epithelial”, “Non-epithelial”) with an expected capture rate of 10,000 cells per library. P-scRNA chambers, one for each population, were each loaded with 30,000 cells with an expected capture rate of approximately 11,000 cells per chamber to approximate similar cell capture in D-scRNA {https://honeycomb.bio/faqs/}, and frozen at -80C for two weeks before library preparation and sequencing. D-scRNA libraries resulted in the capture of more cells (mean = 13,738) than P-scRNA libraries (mean = 4,383) on average, despite the marginally higher expected capture count for P-scRNA. After sample integration and clustering, the largest cell cluster again consisted of cells that could not be assigned to a cell compartment and was primarily composed of P-scRNA cells (**Figure 3A**). In the context of the loaded cell population (epithelial vs. non-epithelial), a predominance of the targeted cell type was observed using both platforms; however, minor proportions of cells corresponding to the non-targeted cell type were also found (i.e., EPCAM transcript-positive cells present in non-epithelial, EPCAM surface marker negative libraries). For P-scRNA libraries, after removing non-typeable and low-quality cells, we assigned a specific type to 2,522 cells for the mixed population, 2,177 cells for the epithelial population, and 402 cells for the “non-epithelial”. In contrast, for D-scRNA libraries, we could assign a specific type for substantially more cells, including 15,120 cells for the mixed population, 4,844 for the epithelial population, and 17,690 for the non-epithelial population. (**Figure 3A and Supplementary Fig. 3**). Overall, a higher fraction (91.4%) of the total D-scRNA cell population from all three D-scRNA libraries could be assigned to a cell type compared to the P-scRNA libraries (38.8%). However, despite the poorer overall capture rates in P-scRNA, most cell types were represented in libraries derived from both platforms, with the notable exception of *OLFM4*-expressing epithelial stem cells, which were surprisingly not found in any of the P-scRNA libraries (**Figures 3A-B)**. Together, these results suggest that D-scRNA methods may capture higher rates of typable cells, particularly rarer cell types, when controlling for the yield of recovered live cells compared to P-scRNA.

**Figure 3.**
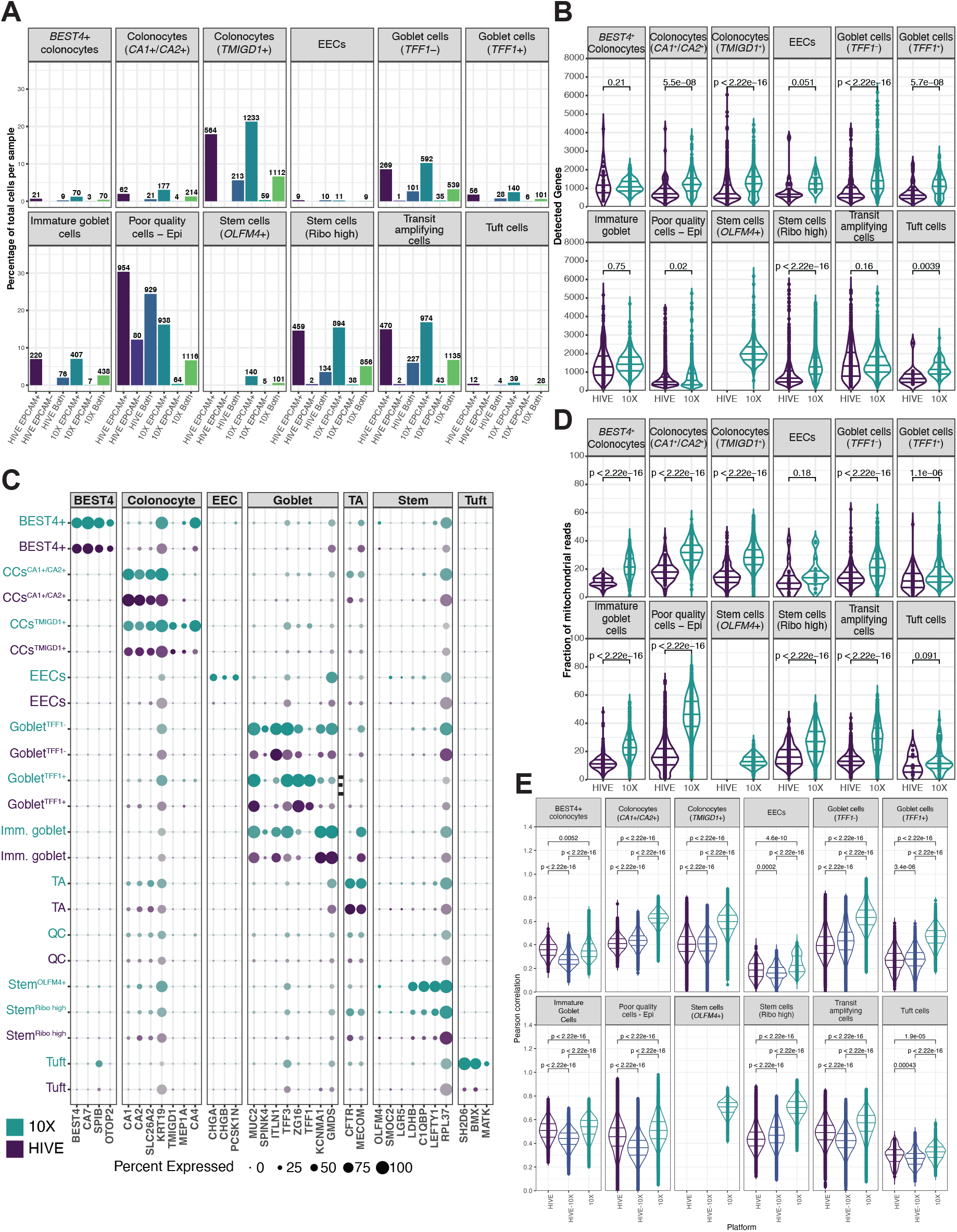
Cell-type specific differences across D-scRNA and P-scRNA platforms (Experiment 3). **A**) The percentage of typeable cell counts per epithelial cell type for each of the P-scRNA and D-scRNA libraries from Experiment 3 with labeled cell count numbers; **B**) the number of detected genes; and **C**) the percentage and expression levels (indicated by color saturation) of known epithelial cell type markers stratified by scRNA platform. **D**) Mitochondrial read fraction per epithelial cell type stratified by scRNA platform. **E**) Distribution of pairwise epithelial cell expression similarity measured by Pearson correlation stratified by scRNA platform. Horizontal lines within violin plots represent the 25^th^, 50^th^, and 75^th^ percentile levels.

To further characterize the performance of the two approaches, we compared the number of UMIs, number of genes, and fraction of mtDNA reads across platforms. P-scRNA libraries had significantly fewer genes and UMIs per cell detected than D-scRNA in most epithelial cell populations (**Figures 3B-C**); however, P-scRNA had higher UMIs and captured genes per cell for most immune and stromal cell types (**Supplementary Figs. 4-5**). These findings are likely a result of the higher number of cells resulting in sequencin*g at less depth in the D-scRNA EPCAM-* and Mixed libraries compared to the D-scRNA libraries in *Experiment 2*, in which fewer cells were captured, showing higher gene and UMIs counts in all compartments compared to P-scRNA. Nonetheless, since individual cell read depth is a function of recovered cells and diluted by the capture of low-quality cells, this result underscores the importance of understanding platform-specific capture rates of the cells being targeted for study during experimental design phases.

**Figure 4:**
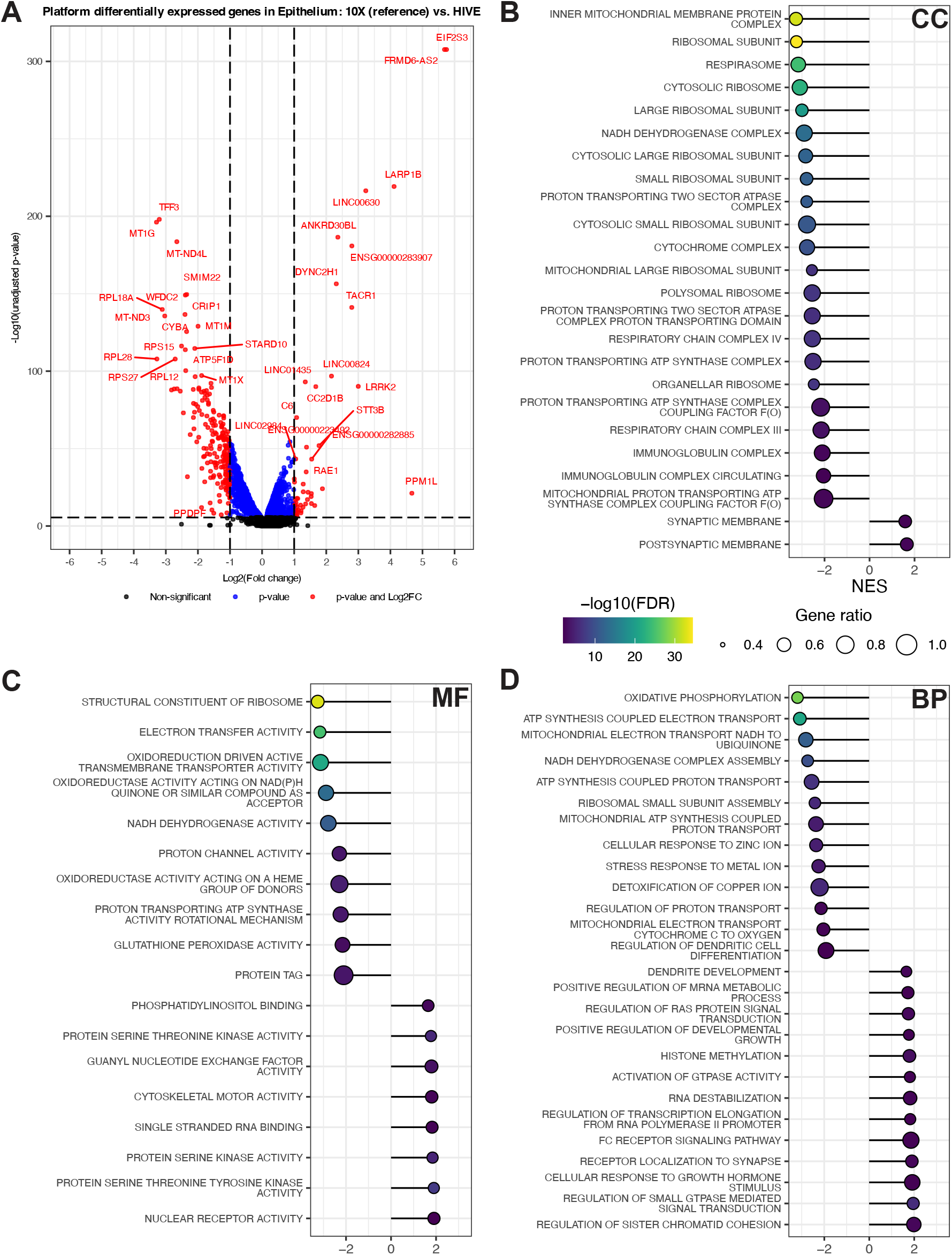
Platform-specific differences observed in genes and gene classes among the same single-cell pools. **A**) Pseudobulk differential expression analysis comparing gene expression of typeable epithelial cells within P-scRNA and D-scRNA libraries included in Experiment 3, restricted to epithelial cells captured in the unsorted and EPCAM+ sorted libraries. Genes to the right are upregulated in P-scRNA libraries relative to D-scRNA while genes to the left are downregulated relative to D-scRNA. The horizontal dotted line indicates a Bonferroni-adjusted p-value of 0.05. **B-D**) Gene Ontology (GO) GSEA of the results from panel A for each GO domain: **B**) cellular component (CC), **C**) molecular function (MF), and **D**) biological process (BP). Normalized expression scores (NES) to the right are enriched terms in P-scRNA libraries while terms to the left are enriched in D-scRNA. Only terms with a p-value <0.01 and gene ratio > 0.50 are shown.

Moreover, platform-specific differences in known cell type markers were found (**Figure 3C and Supplementary Fig. 6**). For example, the proportion of D-scRNA differentiated *TMIGD1*+ colonocytes expressing *CA4*, an accepted marker of colonocytes [8], is significantly higher compared to P-scRNA *TMIGD1*+ colonocytes (83.7% vs 13.6% p-value < 2.2E^-16^; Chi-squared test). Moreover, *TFF3*, a well-described goblet cell marker [9], is significantly differentially expressed compared *TFF3* expression in *TFF1+* to D-scRNA identified goblet cells (**Figure 3C**). However, in most cell types, most notably epithelial cells, the mtDNA read fraction per cell was significantly higher in cells derived from D-scRNA compared to P-scRNA, further supporting that D-scRNA library generation methods stress colonic epithelial cells (**Figure 3D and Supplementary Fig, 7**).

Given the differences in the number of expressed genes per cell between the two platforms, we measured the pairwise Pearson correlation between all cells within each respective cell type to measure how consistently a platform captured the transcriptional profile of a given cell type. D-scRNA cells were significantly more similar to each other (p-value < 0.05, t-test) across almost all cell types compared to P-scRNA cells (**Figure 3E and Supplementary Fig. 8**). To identify potential confounders of cell similarity, we fit multiple univariate linear models using the following measures as predictors of the Pearson correlation: UMI count, expressed gene count, and the percentage difference between P-scRNA and D-scRNA derived cells per cell type. We found that UMI count (R^2^ = 0.265%), expressed gene count (R^2^ = 0.488%), mtDNA read fraction, and the percentage difference between P-scRNA and D-scRNA derived cells per cell type (R^2^ = 4.68%) were weak predictors of cell similarity. Together, these results suggest that D-scRNA captures a cell’s transcriptional profile in the context of cell type more consistently than P-scRNA.

Given the differences in transcriptional profiles identified in the similarity analyses, we next sought to identify whether there are platform-specific biases toward what genes and gene classes are assayed. Across the epithelial compartment, 2,484 genes were significantly differentially expressed (Bonferonni-adjusted p-value < 0.05) between platforms (**Figure 4A**) As highlighted previously, *TFF3* expression is among the top genes where gene expression is significantly lower in P-scRNA compared to D-scRNA in context of cell type. After performing gene set enrichment analysis of the epithelial compartment, we found D-scRNA derived cells tended to be enriched for genes involved in oxidative phosphorylation (likely owing to higher mtDNA fractions) and ribosome function (**Figure 4B-D**). In contrast, P-scRNA cells tended to be enriched for ion channel expression, neurotransmitter receptors, and cytoskeletal proteins. This result highlights that the choice of scRNA platform can affect what types of genes are profiled and that integrating data from different scRNA approaches may provide complementary insights into the transcriptional profile of assayed cells, but specific consideration will be required when considering reproducing biological insights derived from scRNA datasets derived from different technologies.

## Discussion

With the increasing availability of different single-cell RNA expression profiling platforms, our results demonstrate that the choice of platform can have a significant impact on the penultimate data generated based on the advantages and limitations of each approach. In head-to-head comparisons derived from the same single-cell pools, we find that D-scRNA methods may allow for the capture of more cells overall, including a more faithful representation of rare cell types, with higher and more consistent gene expression in the context of cell type. On the other hand, P-scRNA approaches that allow for cryopreservation and broader in-field applications may result in the capture of fewer high-quality cells, highlighting the need for careful quality control measures before downstream analyses. Our comparisons also highlight clear platform differences in the types of transcripts captured, suggesting downstream analyses from merged datasets across platforms may be challenging without additional experimentation and bridging controls to adjust for technical biases. Further, cell-specific markers may differ between the two platforms which highlights the need to exercise additional caution when comparing or contrasting results across platforms, as observed for *TFF3* expression in *TFF1*+ goblet cells. Given that many of the routinely used colonic mucosa cell type markers have been determined from 10X Chromium, additional effort may be required to ensure the translatability of these markers to alternative platforms.

Notably, since this study was performed, Honeycomb has released HIVE CLX, the next generation of their P-scRNA platform, which is reported to capture 4x more cells and 20% more genes/UMIs. Despite these improvements, our results suggest that the cell capture rate will likely still fall short of the D-scRNA overall capture rates of high-quality cells from loaded cells for human colon biopsies. However, this advantage of D-scRNA may be derived at the expense of higher cell injury/stress rates, as evidenced by higher mtDNA read fractions than P-scRNA, particularly among epithelial cell types. Therefore, although D-scRNA may capture more cells than P-scRNA, cellular injury induced by the library preparation methods for D-scRNA may alter the native transcriptional profile of epithelial cells and subsequent analyses will need to carefully control for these potential effects.

Our work also highlights the challenges of profiling human mucosal biopsies at single-cell resolution and the need for innovation to improve platforms that are more conducive to translational research. The most well-established platforms require samples to be collected from the patient and immediately brought to a well-equipped lab to be dissociated and sorted, regardless of when samples may ultimately undergo library preparation (i.e., immediately, as is the case for D-scRNA methods or cryopreserved for later preparation as is the case for P-scRNA). Moreover, in the case of D-scRNA, which appears particularly harsh on epithelial cells, careful attention to optimizing collection and library preparation steps is required to avoid significant sample quality degradation. To address poor capture of fragile cell types in fresh samples, 10X Genomics has developed the “Flex” assay, which uses formaldehyde to fix cellular RNA prior to D-scRNA sequencing. This approach could potentially address the damaging effects of D-scRNA sequencing on colonic epithelial cells. Alternatively, the P-scRNA approach does allow storage for up to 2 weeks before library preparation which appears to perform almost as well as samples freshly prepared using this methodology. This not only allows stable transport of processed cells to other sites equipped for library preparation and sequencing but also joint library preparation across samples and sequencing, potentially limiting potential batch effects when compared to D-scRNA approaches. Notably, in our hands, we found that the -20C storage condition recommended by the manufacturer was inadequate to ensure viable cell recovery of cells prepared from human rectosigmoid colon biopsy samples but found that -80C samples captured cells similarly to cells processed immediately without cryopreservation.

Taken together, our results provide important methodological context for the interpretation of both scRNA approaches and highlight additional areas needing improvement to broaden the adoption of these technologies in large-scale human studies, particularly those using intestinal biopsies or focusing on epithelial cell biology. Nonetheless, within the context of these limitations, deriving high-quality single-cell data is still possible, but additional emphasis must be placed on optimizing sample collection, bioinformatic pipelines, and analyses for specific platforms, as our results highlight that not all methods are broadly similar. This has obvious repercussions when considering rigor and reproducibility across studies that employ different platforms while trying to interrogate similar cellular processes within the same tissue and even cell types. These results also highlight the importance of advancing methods to allow for single-cell transcriptomics in fixed or otherwise preserved tissues to provide additional opportunities for validation. Such progress would also overcome the challenges of acquiring fresh tissue and requiring immediate processing, which is time-sensitive, laborious, and resource-intensive.

## Methods

All subjects provided informed written consent for the study prior to participating. The study protocol was approved by the Dana-Farber/Harvard Cancer Center Institutional Review Board (Boston, MA, DF/HCC #21-376; NCT05056896). Six jumbo pinch biopsies of grossly normal-appearing mucosa were taken via flexible sigmoidoscopy in the rectosigmoid colon (15-20 cm from the anal verge). The biopsies were immediately placed into a 1.5 mL cryovial with collection media (79% ADMEM/F12K, 20% sterile and filtered FBS, and 1% L-POPS) on wet ice. This media was selected on the basis that human colon biopsies collected under this condition for organoid cultures, which require viable epithelial cells, routinely yield successful derivation after short-term storage (<1 hour) on wet ice during transport from clinical procedures to a processing laboratory [10]. Biopsies were mechanically disrupted and digested with dithiothreitol (DTT)/collagenase/Dnase immediately within 30 minutes of collection. Suspended cells were then FACS sorted for viability and sorted into epithelial (EPCAM+/CD45-) and non-epithelial fractions (EPCAM-), where applicable. HIVE libraries were created using the Honeycomb HIVE v1 system with the guidance of the manufacturer’s published protocol. The 10X Chromium libraries were generated using the 3’ v3.1 chemistry after loading a volume containing at maximum 10,000 cells onto a single 10X Chromium lane. Libraries from both platforms were sequenced on the Illumina NovaSeq platform to a depth of 100-200M reads per sample to the manufacturer’s specified length.

We used the *alevin-fry*/*simpleaf* (version 0.14.1) pipeline to perform read mapping, cell-barcode correction, empty-droplet removal, and per-cell gene count generation [11, 12] to avoid confounding introduced by the differences in read mapping and gene counts-by-cell matrix generation. Intronic, exonic, and ambiguous genomic reads were aligned to the GRCh38 genome and the GENCODE v44 transcript list. Given the difference in read length between the HIVE and 10X platforms, separate reference genome indices were generated and used for alignment. The HIVE cell barcode length on R1 reads was set to 12 base pairs (bp) followed by 13 bp of random UMI sequence. The 10X R1 reads followed v3 chemistry standards. The HIVE reference index was generated using an R2 read length of 51 with k-mer sizes of 25, whereas the 10x index assumed R2 read lengths of 90 with k-mer sizes of 31. The removal of empty droplets was performed using the “knee” method without using 10x’s cell barcode whitelist, as HIVE does not have such a list.

*Seurat* v5 performed per-cell quality control and downstream analyses within the R (v4.3.1) environment [13]. Cells with a gene count < 200 or were found to be a doublet by *scDblFinder* [14] were removed. Cells with high mitochondrial read fractions were not removed from the analysis to highlight storage and platform differences in epithelial cell injury during sample processing. Gene counts were normalized using the Seurat *SCtransform* function [15] while regressing out the effects of mitochondrial read fraction. Sample integration was performed using Harmony [16] using 50 principal components to control for the effects of library preparation and cell sorting methods. Cell typing was first performed by subdividing cell clusters into epithelial, stromal, and immune compartments, followed by inspecting the expression patterns of known cell-specific markers [2, 3, 17-19]. Cell clusters defined by mitochondrial genes, lncRNAs (such as *MALAT1*), genes not native to the cell compartment (e.x. Immune gene expression in epithelial cells), or lack of specific genes to enable cell typing were defined as low-quality cells. Pairwise cell similarity was performed by calculating the Pearson correlation between the normalized expression values of each cell type’s 2,000 most variable genes. *DESeq2* was used to identify scRNA platform-associated differentially expressed genes within the epithelial compartment [20]. Specifically, all gene expression counts from quality epithelial cells were aggregated in a library-specific manner (“pseudobulked”). Only the EPCAM+ sorted and unsorted P-scRNA and D-scRNA libraries were considered for differential gene expression analysis given the low number of epithelial cell counts in the EPCAM-libraries. Gene Ontology (GO) [21, 22] gene set enrichment analysis (GSEA) was performed by *clusterProfiler* [23].

## Abbreviations

D-scRNA: droplet-based scRNA
GO: Gene Ontology
GSEA: gene set enrichment analysis
P-scRNA: picowell-based scRNA
scRNA: single-cell RNA sequencing
UMI: unique molecular identifier

## Author Contributions

Conceptualization: JMD, CMG, OHY, TS, ATC, DAD; Data curation: JMD, RJM; Formal Analysis: JMD, RJM, ATC, DAD; Funding acquisition: OHY, TS, ATC, DAD; Investigation: JMD; OHY, TS, ATC, DAD; Methodology: JMD, CMG, DAD; Resources: OHY, TS, ATC, DAD; Supervision: OHY, TS, ATC, DAD; Visualization: JMD; Writing – original draft: JMD, DAD; Writing – review & editing; All authors.

## Acknowledgments

We would like to acknowledge the following members of the Massachusetts General Hospital clinical research coordination team, who were critical in enrolling the participants of this study, collecting biospecimens, and maintaining scientific rigor for generation of these data: Marina Magicheva-Gupta, Ingrid Crumpton, Liza Hote, Karen Kirunda, Madeline Koehn, Perry Labelle, Trenton Reinicke, Janavi Sethurathnam, and Julia Solowey as well as all clinical investigators on the ASPIRED-XT trial who assist with clinical procedures, and individuals who participated on the ASPIRED-XT trial.

