## Supplementary Figures for "Optimizing single-cell RNA sequencing methods for human colon biopsies: droplet-based vs. picowell-based platforms"

Supplementary Figure 1. UMAP plots and gene markers plots from experiment 1 A) UMAP of HIVE Fresh vs. -80C comparison; all cells B) Dot plot of cell type markers within the two HIVE libraries. Nontypeable cells showed low expression of markers found in all three cell compartments. C) UMAP of HIVE Fresh vs. -80C comparison; epithelial cells. D) UMAP of HIVE Fresh vs. -80C comparison; immune cells. E) UMAP of HIVE Fresh vs. -80C comparison; stroma cells.

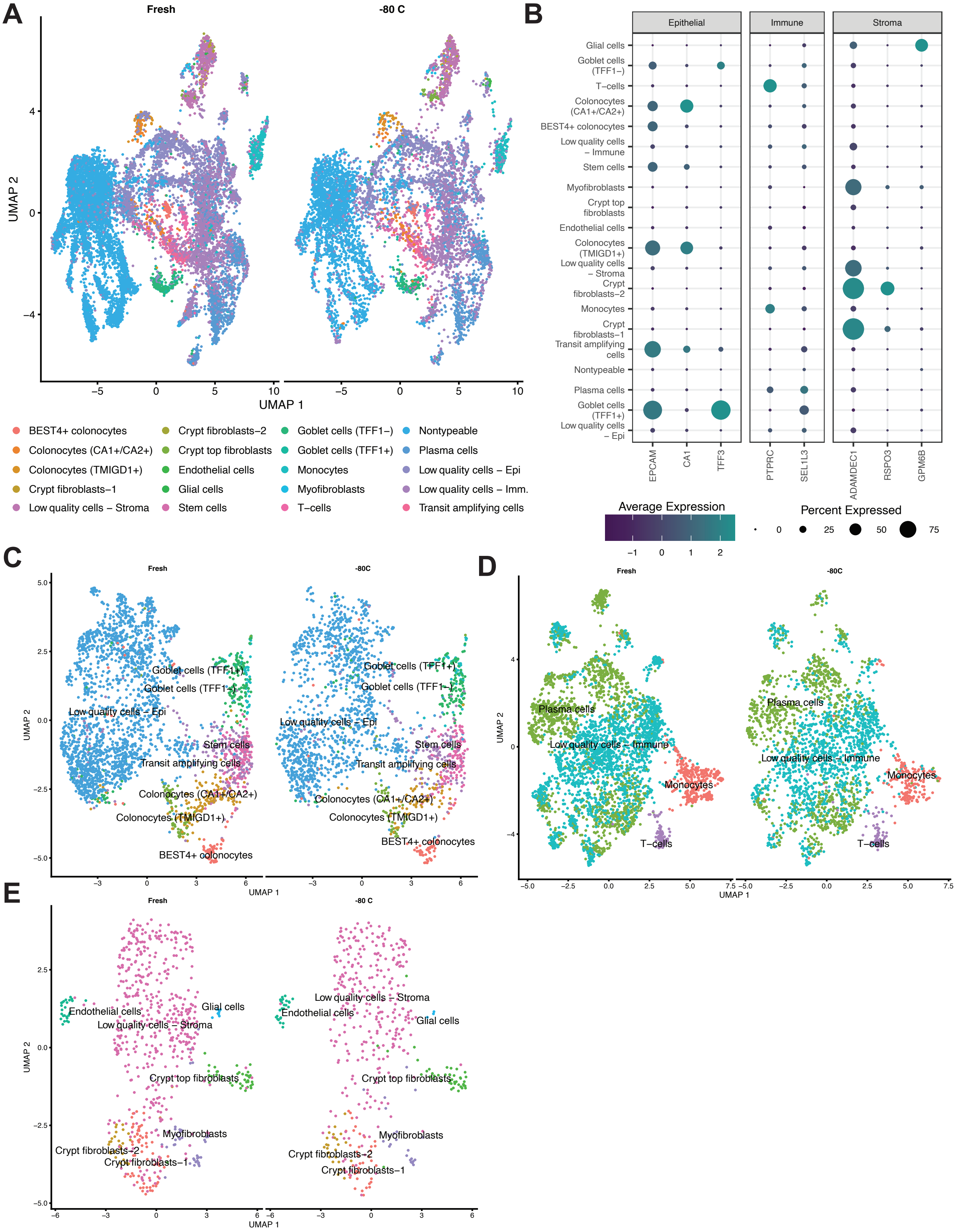

Supplementary Figure 2. A) Percentage of total cell counts per cell type across all mucosal compartments for the HIVE Fresh/-80 to 10X comparison with labeled cell count numbers after removing nontypeable cells. B) The number of detected genes per cell type across all mucosal compartments for the HIVE Fresh/-80 to 10X comparison. C) Unique molecular identifiers (UMI per cell type across all mucosal compartments for the HIVE Fresh/-80 to 10X comparison. D) Mitochondrial read fraction per cell type across all mucosal compartments for the HIVE Fresh/-80 to 10X comparison.

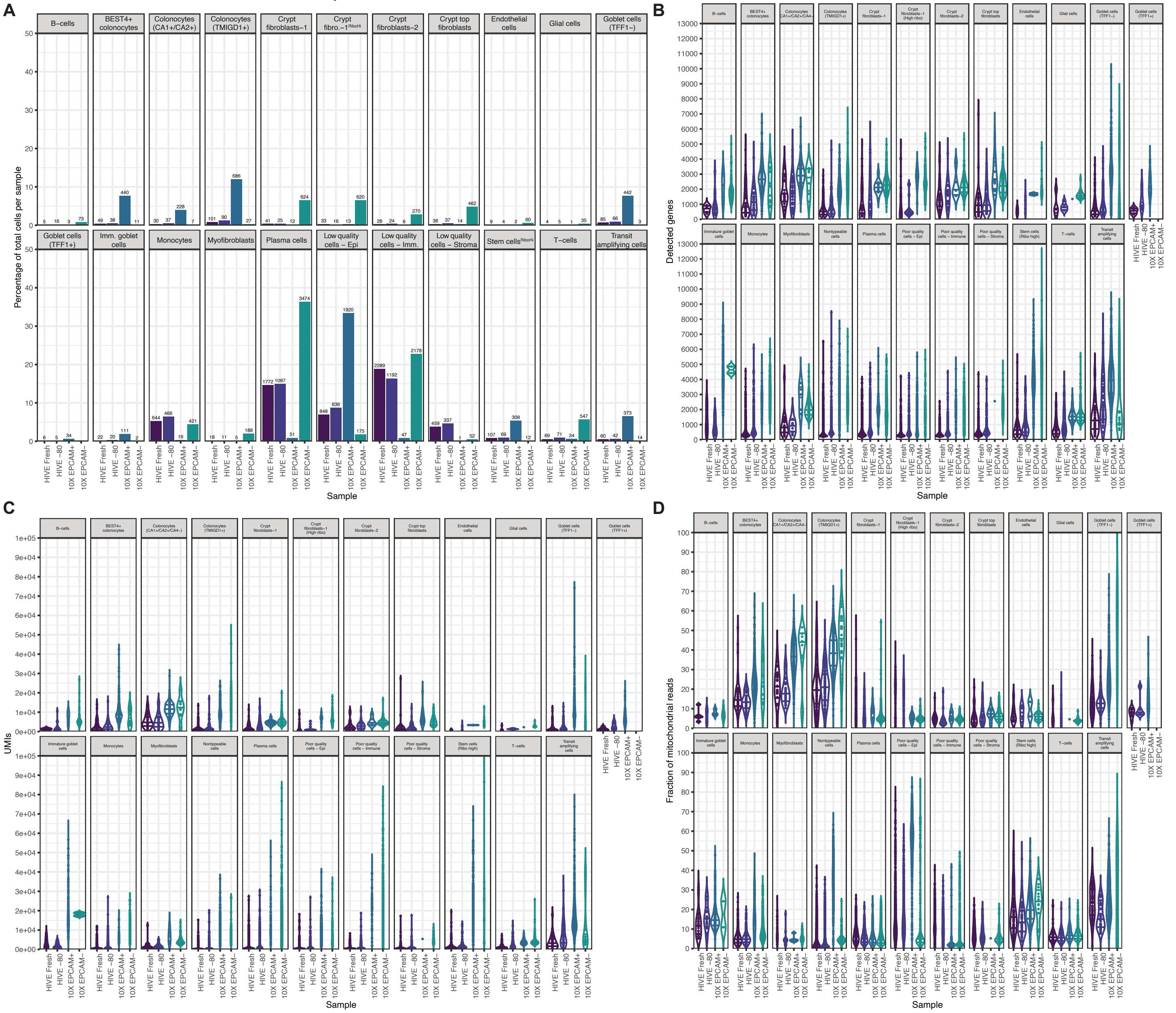

Supplementary Figure 3: A) UMAP of HIVE and 10X Chromium libraries in Experiment 3; all cells. B) The percentage of total cell counts per cell type for each of the HIVE and 10X libraries from Experiment 3 with labeled cell count numbers after the removal of nontypeable cells – Epithelium. C) The percentage of total cell counts per cell type for each of the HIVE and 10X libraries from Experiment 3 with labeled cell count numbers after the removal of nontypeable cells – Immune. D) The percentage of total cell counts per cell type for each of the HIVE and 10X libraries from Experiment 3 with labeled cell count numbers after the removal of nontypeable cells – Stroma.

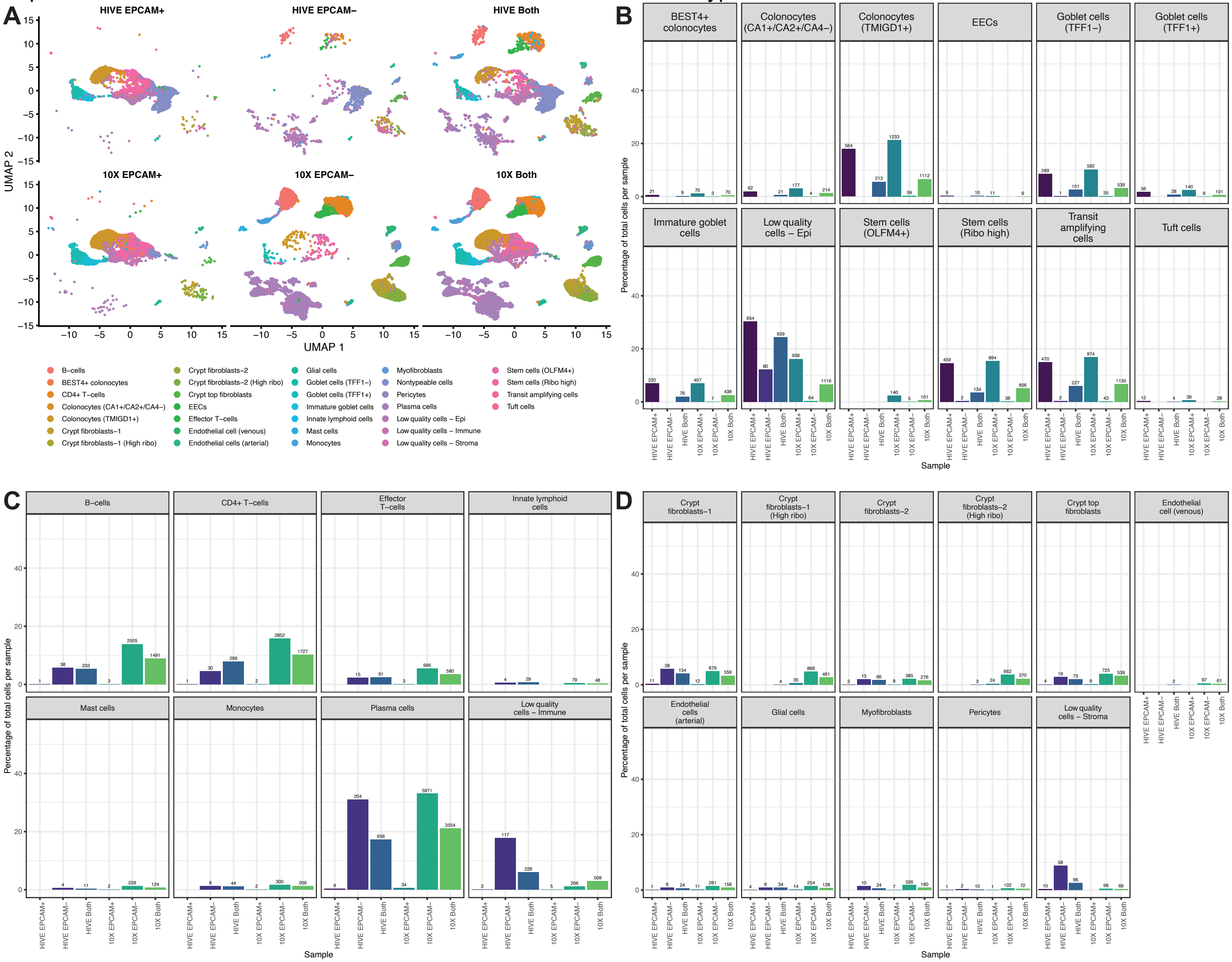

Supplementary Figure 4: A) The number of detected genes per cell type stratified by scRNA platform used in Experiment 3 (10X vs HIVE) – Immune. B) The number of detected genes per cell type stratified by scRNA platform used in Experiment 3 (10X vs HIVE) – Stroma.

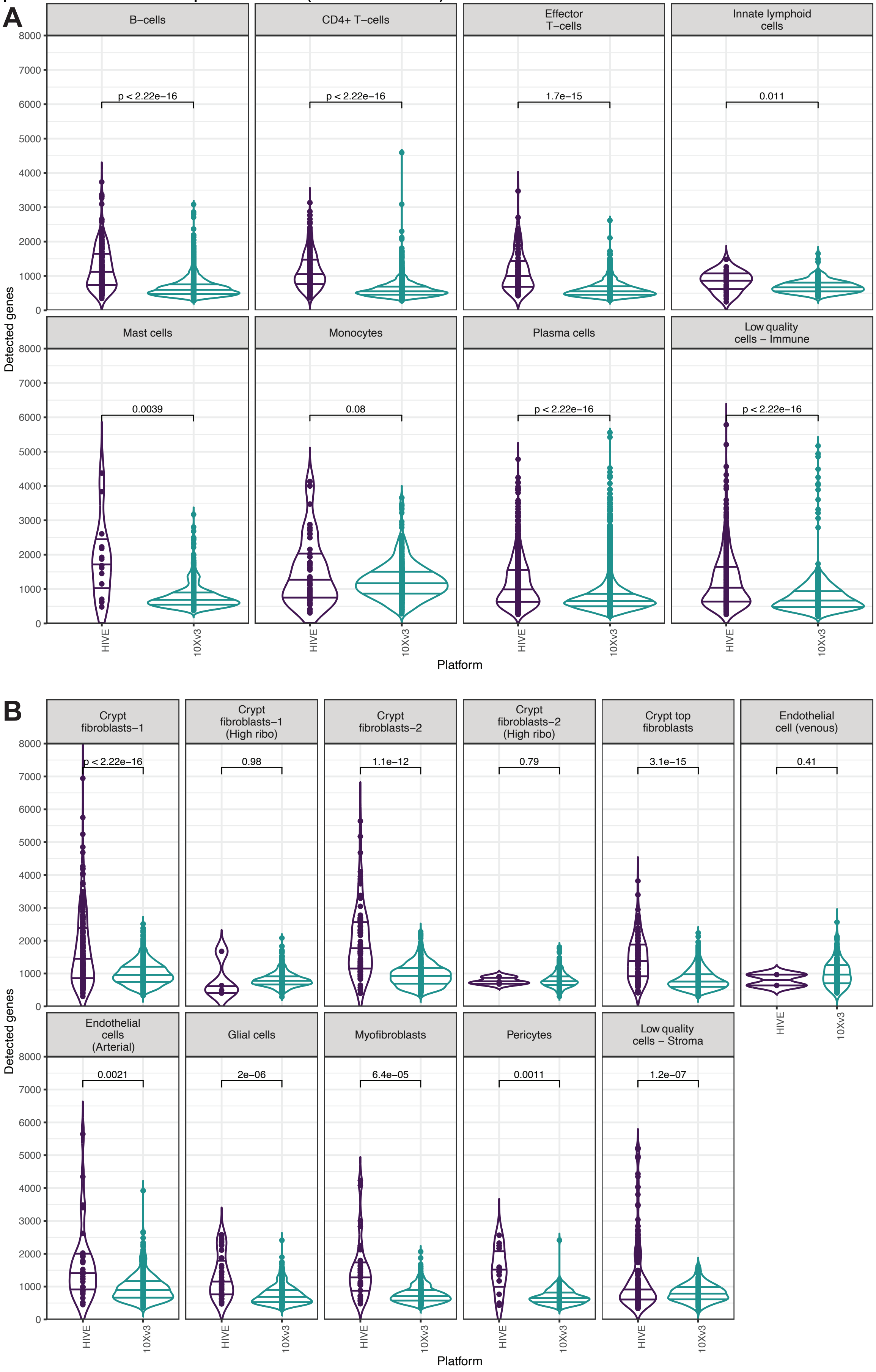

Supplementary Figure 5: A) The number of UMIs per cell type stratified by scRNA platform used in Experiment 3 (10X vs HIVE) – Epithelium. B) The number of UMIs per cell type stratified by scRNA platform used in Experiment 3 (10X vs HIVE) – Immune. C)

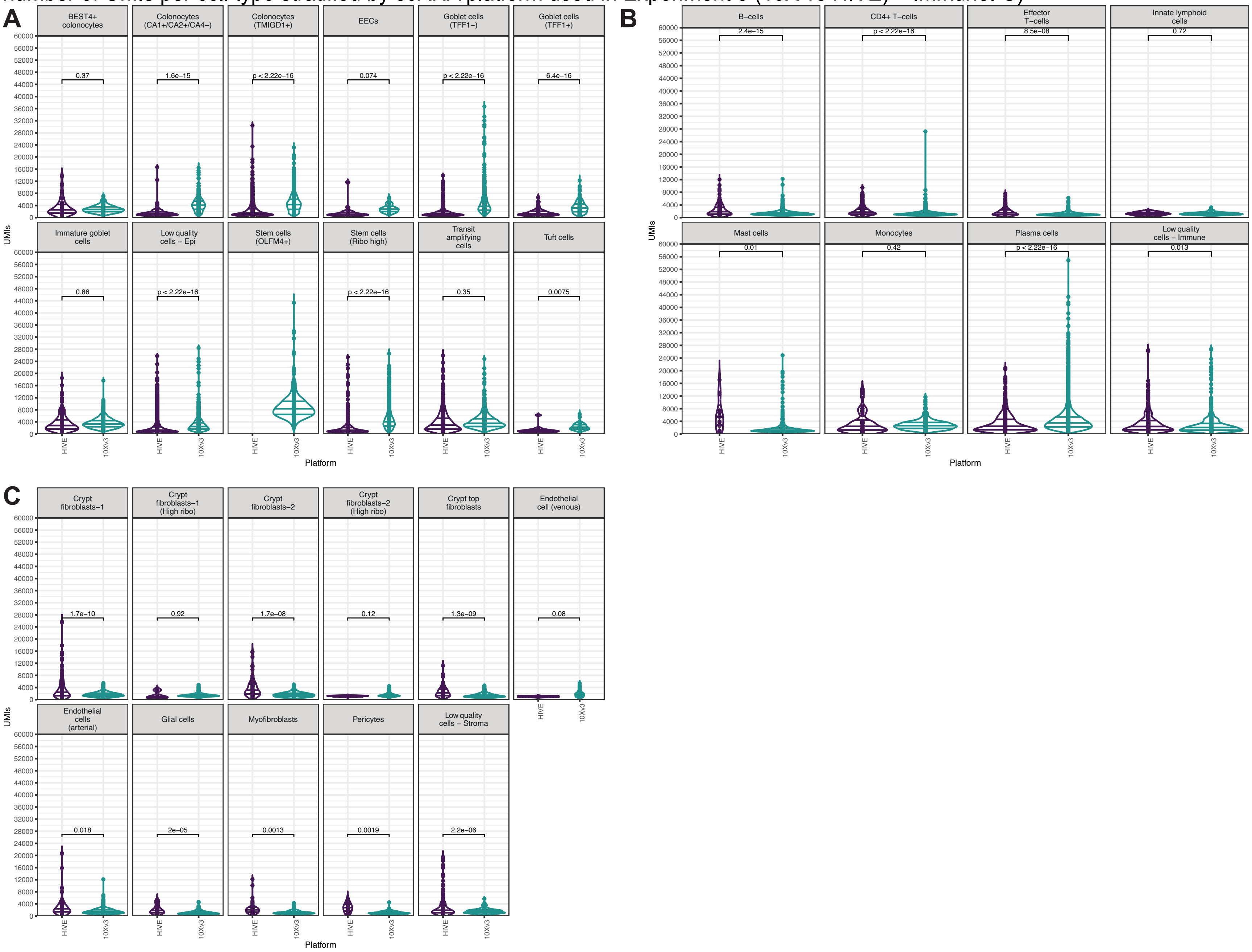

Supplementary Figure 6: A) The percentage and expression levels (indicated by color saturation) of known cell type markers stratified by scRNA platform – Immune. B) The percentage and expression levels (indicated by color saturation) of known cell type markers stratified by scRNA platform – Stroma

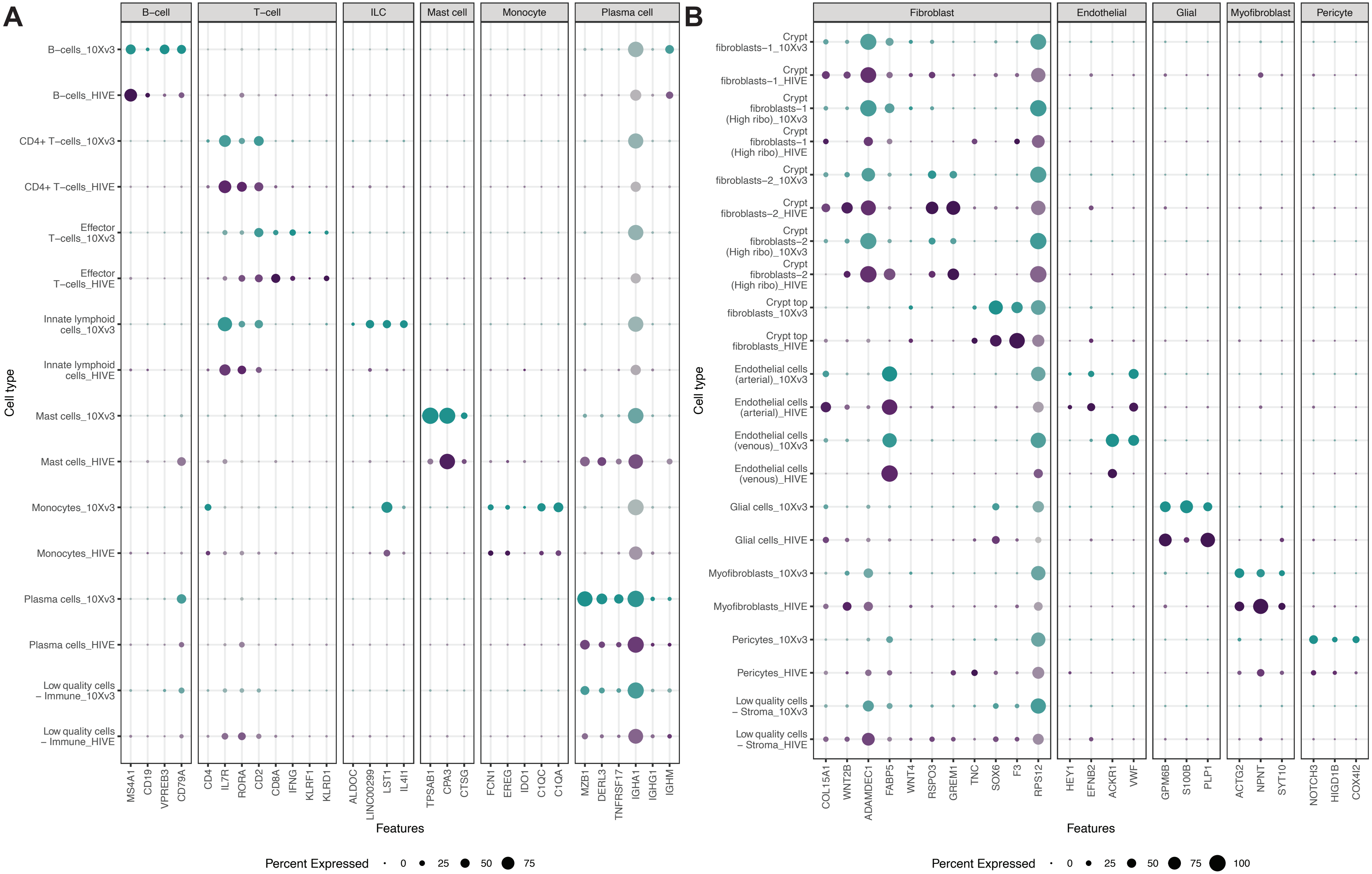

Supplementary Figure 7: A) The fraction of mitochondrial reads per cell type stratified by scRNA platform used in Experiment 3 (10X vs HIVE) – Immune. B) The fraction of mitochondrial reads per cell type stratified by scRNA platform used in Experiment 3 (10X vs HIVE) – Stroma

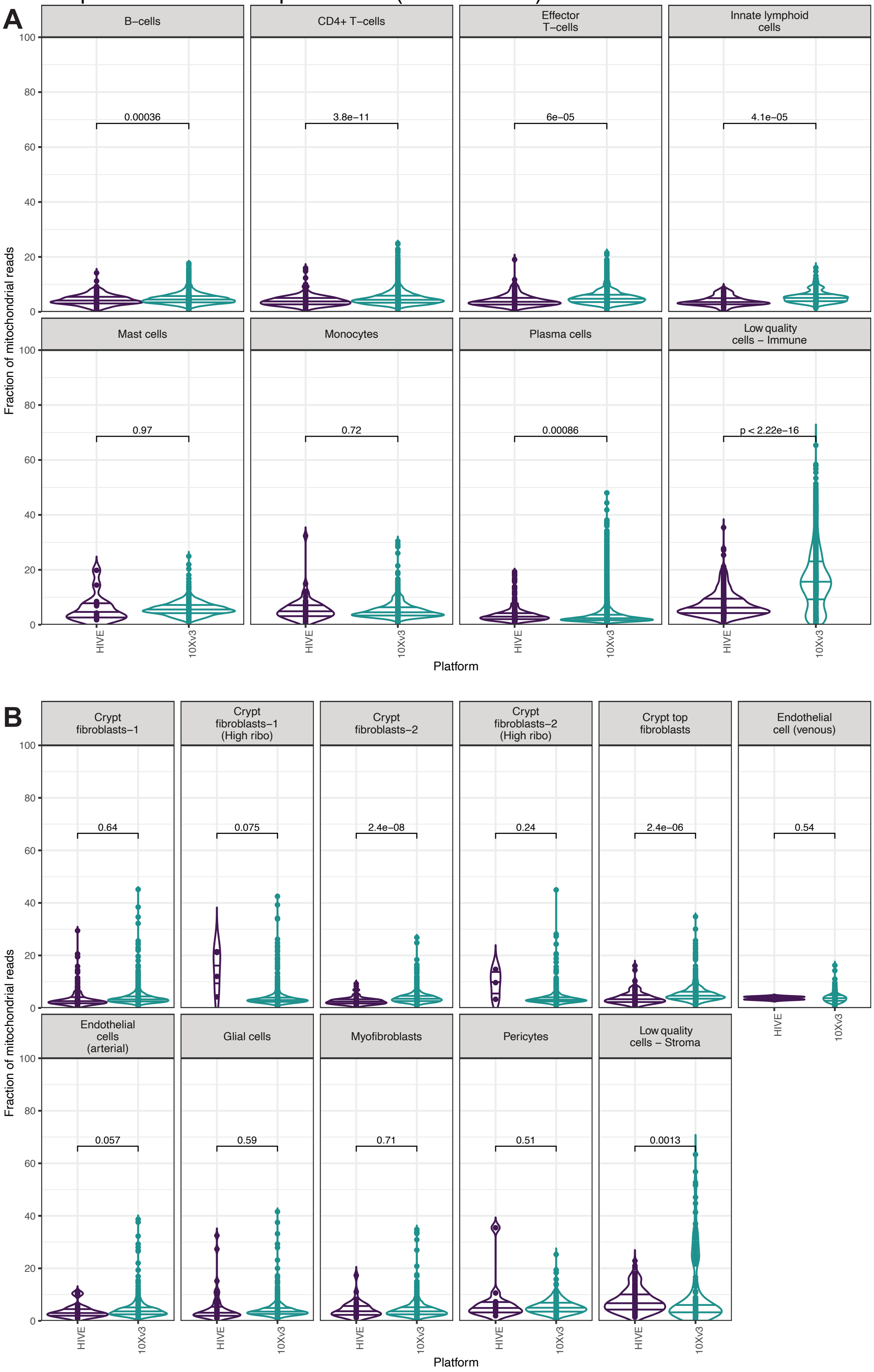

Supplementary Figure 8: A) The distribution of pairwise cell expression similarity measured by Pearson correlation stratified by scRNA platform – Immune. B) The distribution of pairwise cell expression similarity measured by Pearson correlation stratified by scRNA platform – Stroma

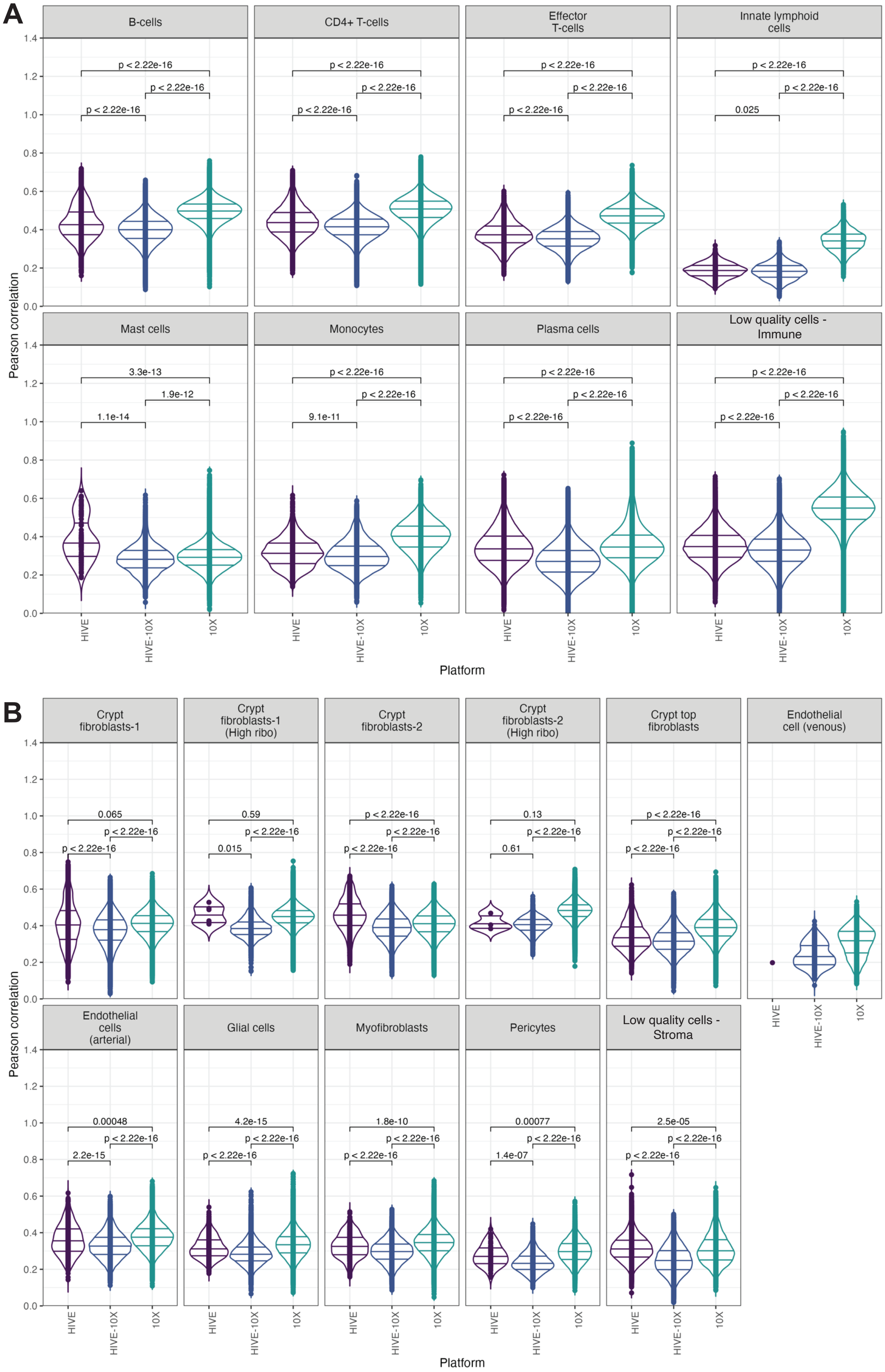
